# To denoise or to cluster? That is not the question. Optimizing pipelines for COI metabarcoding and metaphylogeography

**DOI:** 10.1101/2021.01.08.425760

**Authors:** A. Antich, C. Palacin, O.S. Wangensteen, X. Turon

**Affiliations:** Department of Marine Ecology, Centre for Advanced Studies of Blanes (CEAB-CSIC), Blanes (Girona), Catalonia, Spain; Department of Evolutionary Biology, Ecology and Environmental Sciences, University of Barcelona, and Research Institute of Biodiversity (IRBIO), Barcelona, Catalonia, Spain; Norwegian College of Fishery Science, UiT The Arctic University of Norway, Tromsø, Norway

## Abstract

The recent blooming of metabarcoding applications to biodiversity studies comes with some relevant methodological debates. One such issue concerns the treatment of reads by denoising or by clustering methods, which have been wrongly presented as alternatives. It has also been suggested that denoised sequence variants should replace clusters as the basic unit of metabarcoding analyses, missing the fact that sequence clusters are a proxy for species-level entities, the basic unit in biodiversity studies. We argue here that methods developed and tested for ribosomal markers have been uncritically applied to highly variable markers such as cytochrome oxidase I (COI) without conceptual or operational (e.g., parameter setting) adjustment. COI has a naturally high intraspecies variability that should be assessed and reported, as it is a source of highly valuable information. We contend that denoising and clustering are not alternatives. Rather, they are complementary and both should be used together in COI metabarcoding pipelines. Using a typical dataset from benthic marine communities, we compared two denoising procedures (based on the UNOISE3 and the DADA2 algorithms), set suitable parameters for denoising and clustering COI datasets, and compared the outcome of applying these processes in different orders. Our results indicate that denoising based on the UNOISE3 algorithm preserves a higher intra-cluster variability. We suggest and test ways to improve this algorithm taking into account the natural variability of each codon position in coding genes. The order of the steps (denoising and clustering) has little influence on the final outcome. We recommend researchers to consider reporting their results in terms of both denoised sequences (a proxy for haplotypes) and clusters formed (a proxy for species), and to avoid collapsing the sequences of the latter into a single representative. This will allow studies at the cluster (ideally equating species-level diversity) and at the intra-cluster level, and will ease additivity and comparability between studies.

## INTRODUCTION

The field of eukaryotic metabarcoding is witnessing an exponential growth, both in the number of communities and substrates studied and the applications reported (reviewed in Deiner et al 2017, Aylagas et al 2018, Bani et al 2020, Compson et al 2020). In parallel, technical and conceptual issues are being discussed (e.g., Mathieu et al 2020, Rodriguez-Ezpeleta et al 2020) and new methods and pipelines generated. In some cases, however, new practices are established after a paper reporting a technique is published and followed, sometimes pushing its application outside the context in which it was first developed.

A recently debated matter concerns the treatment of reads by denoising procedures or by clustering techniques (Porter & Hajibabaei 2020). Both methods are often presented as alternative approaches to the same process (e.g., Macheriotou et al 2018, Forster et al 2019, O’Rourke et al 2020, Giebner et al 2020). However, both are philosophically and analytically different (Turon et al 2020). While denoising strives to detect erroneous sequences and to merge them with the correct “mother” sequence, clustering tries to combine a set of sequences (without regard to whether they contain or not errors) into meaningful biological entities, ideally approaching the species level, called OTUs or MOTUs (for Molecular Operational Taxonomic Units). Usually only one representative sequence from each MOTU is kept (but note that this is only common practice, not a necessary characteristic of the method). Thus, while both procedures result in a reduced dataset and in error correction (by merging reads of erroneous sequences with the correct one or by combining them with the other reads in the MOTU), they are not equivalent. More importantly, they are not incompatible at all and can (and should) be used together.

A recent paper by Callahan et al (2017) suggests that denoised sequences should replace MOTUs as the unit of metabarcoding analyses. We contend that it may be so for ribosomal DNA datasets such as the one used in that paper, but that this notion has gained momentum also in other fields of metabarcoding for which it is not adequate. In particular, when it comes to highly variable markers such as COI. The 3’ half (also called Leray fragment) of the standard barcode fragment of COI (Folmer fragment) is becoming a popular choice for metabarcoding studies addressed at metazoans or at eukaryotic communities at large (Andújar et al 2018), reaching now 28% of all metabarcoding studies (van der Loos & Nijland 2020). Metabarcoding stems from studies of microbes where 16S rRNA is the gene of choice, and the concept was then applied to 18S rRNA analyses of eukaryotes. With the recent rise of COI applications in metabarcoding, programs and techniques developed for rDNA are sometimes applied to COI without reanalysis and with no parameter adjusting given the highly contrasting levels of variation of these markers.

The idea that denoising should be used instead of clustering has been followed by some (e.g., Tapolczai et al 2019, Holman et al 2020, Steyaert et al 2020, Zamora-Terol et al 2020, Pearman et al 2020), while other authors have combined the two approaches (e.g., Brandt et al 2020, Nguyen et al 2020, Laroche et al 2020). Indeed, denoising has the advantages of reducing the dataset and to ease pooling or comparing studies, which is necessary in long term biomonitoring applications. However, with COI there is a wealth of intraspecific information that can be used and that is lost if only denoising is applied (Zizka et al 2020). Not in vain COI has been a prime marker of phylogeographic studies to date (Avise 2009, Emerson et al 2011), and these studies can be extended to metabarcoding datasets by mining the distribution of haplotypes within MOTUs (metaphylogeography, Turon et al 2020). The latter authors suggested to perform clustering first, and that denoising should be done within MOTUs to provide the right context of sequence variation and abundance skew. They also advised to perform a final abundance filtering step. In other studies, denoising is performed first, followed by clustering and refining steps (e.g., Nguyen et al 2020, Laroche et al 2020).

There are several methods for denoising (reviewed in Peng & Dorman 2020) and for clustering (reviewed in Kopylova et al 2016). We will use two of the most popular denoising techniques, based on the DADA2 algorithm (Divisive Amplicon Denoising Algorithm, Callahan et al 2016) and the UNOISE3 algorithm (Edgar 2016). The results of the former are called Amplicon Sequence Variants (ASVs) and those of the latter ZOTUs (zero-radius OTUs). In practice, the terminology is mixed and ASV, ZOTU, ESV (Exact Sequence Variant), sOTU (sub-OTU) or ISU (Individual Sequence Variant), among others, are used more or less interchangeably. For simplicity, as all of them are equivalent, we will use henceforth the term ESV. Clustering, on the other hand, can be performed using similarity thresholds (e.g., Edgar 2013, Rognes et al 2016), Bayesian Methods (CROP, Hao et al 2011), or methods based on single-linkage-clustering (SWARM, Mahé et al 2015), among others. We will focus on *de novo* clustering methods (i.e., independent of a reference database), while denoising is always *de novo* by its very nature (Callahan et al 2017). We will here use SWARM as our choice of clustering program due to its good performance compared to other methods (Kopylova et al 2016). It is noteworthy that all these programs were originally developed and tested on 16S rDNA datasets. When applied to other markers, often no indication of parameter setting is given (i.e., omega_A for DADA2, α for UNOISE3, d for SWARM), suggesting that default parameter values are used uncritically.

In this article, we aim to use a COI metabarcoding dataset of benthic littoral communities to (1) set the optimal parameters of the denoising and clustering programs for COI markers, (2) compare results of the DADA2 algorithm with the UNOISE3 algorithm, (3) compare the results of performing only denoising, only clustering, or combining denoising with clustering in different orders, and (4), suggest and test improvements in the preferred denoising algorithm to take into account the fact that COI is a coding gene. Our aims are to provide guidelines for using these key bioinformatic steps in COI metabarcoding and metaphylogeography.

## MATERIAL AND METHODS

### The dataset

We used as a case study an unpublished dataset of COI sequences obtained from benthic communities in 12 locations of the Iberian Mediterranean. The seaweed-dominated shallow community inhabiting vertical rocky surfaces between −4 and −8 m was sampled by completely scraping off with hammer and chisel standardized surfaces of 25*25 cm. Three replicate samples were taken per location, and all samplings were performed in autumn of 2017. Sampling localities and coordinates are given in Figure S1.

Sample processing was based in Wangensteen et al (2018) and included a size fractionation step. Extraction and amplification was also performed as in that work using a modified version of the Leray et al (2013) primer set (called Leray-XT in Wangensteen et al 2018), which also added unique 8-bp sample tags at both ends. HTS library preparation was performed using the NextFlex PCR-free DNA-Seq kit (Perkin-Elmer), based on ligation of the Illumina adapters at both ends of the amplicons. See below for the implications of PCR-free library construction methods in the application of one of the denoising algorithms (DADA2). We used a full run of a V3 Illumina MiSeq kit with 2*250 bp paired-end sequencing.

### Bioinformatic analyses

The initial steps of the bioinformatic pipeline followed Turon et al (2020) and were based on the OBItools package (Boyer et al 2016). Reads were paired and quality filtered, demultiplexed, and dereplicated. A strict length filter of 313 bp was used. We also eliminated sequences with only one read. Chimera detection was performed on the whole dereplicated dataset with uchime3_denovo as embedded in unoise3 (USEARCH 32-bit free version, Edgar 2010). We used minsize=2 to include all sequences. Those identified as chimeras were recovered from the –tabbedout file and eliminated from the dataset. Sequences with small offsets (misaligned), identified as shifted in the output, were likewise deleted. The working dataset thus comprised well-aligned, chimera-free, unique sequences which had appeared with at least two reads in the samples.

Note that for this technical study we didn’t take into account the sample distribution of the reads. A complete biogeographic study of the samples is ongoing and will be published elsewhere. For the present analysis, for each unique sequence only the actual DNA sequence and the total number of reads were retained.

### The denoisers: UNOISE3 and DADA2

Comparing denoising algorithms is challenging because each method comes with a different software suite with embedded features and recommendations (Peng & Dorman 2020). For instance, uchime3_de novo is embedded in the unoise3 command as implemented in USEARCH, while a chimera removal procedure (removeBimeraDenovo) is an optional feature in the DADA2 pipeline. Furthermore, while UNOISE3 uses merged reads, DADA2 recommends denoising forward and reverse reads separately, and then performing a merging step. We have tried to isolate the algorithms from their pipelines for comparability. This was done by generating a Python script that implements the algorithm described in Edgar (2016) and by using DADA2 from its R package v. 1.14.1 and not as embedded into the qiime2 pipeline (Bolyen et al 2019), and then altering some of the recommended procedures and testing different values for the stringency parameters.

For UNOISE3, our program (henceforth DnoisE) was compared on the working dataset described above with command unoise3 in USEARCH with minsize=2, alpha=5 and without the otutab step. That is, we recovered the ESV composition and abundance with an R script directly from the output of unoise3 (using the output files –tabbedout and –ampout), without a posterior re-assignment of sequences to ESVs via otutab. This step was not necessary as all sequences were included in the ESV calculations. The results of DnoisE and unoise3 were >99.99 % identical in ESVs recovered and reads assigned to them, so we continued to use our script for performing the comparisons and for further improvements of the algorithm (see below).

The recommended approach for DADA2 is to denoise separately the forward and reverse reads of each sequence. This complicates the technical comparison, as all initial filtering steps cannot be equally performed (f.i., we won’t know if there is just one read of a particular sequence, or if the merged pair will be discarded for low quality of the assembly or for unsuitable final length) and thus we cannot have two identical starting datasets. More importantly, we cannot use this procedure when we test the effects of denoising at later steps (i.e., after clustering), so we would be unable to compare the denoisers at this level. Thus, for our comparative analysis we need to use DADA2 on merged reads. According to Callahan et al (2016), this can result in a loss of accuracy, but this point has never been tested to our knowledge. We addressed this issue by comparing denoising before and after pairing on half of the reads in the final dataset. After this analysis, we decided to continue our comparison of DADA2 and UNOISE3 on paired reads.

Additionally, using reads before pairing is not optimal if a PCR-free library preparation protocol is used, as in our case, because half of the reads are in one direction and the other half are in the opposite direction (hence the use of half of the reads in the above comparison). Forward and reverse reads can of course be recombined to generate new files with all reads in the same direction, but the quality of the original forward and reverse reads is different. Alternatively, two rounds of DADA2 (one per orientation) must be performed and combined at later steps.

To run DADA2 on merged reads, we entered them in the program as if they were the forward reads and did not use a merging step after denoising. In all DADA2 runs we did not perform the recommended chimera removal procedure as the input sequences were already chimera-free according to uchime3_de novo. Note that, when denoising was done after clustering, we used error rates calculated for the whole dataset, and not for each MOTU separately (most of them do not have enough number of sequences for a reliable estimation of error rates).

UNOISE3 relies heavily on the stringency parameter α. In short, lower values of α tend to merge sequences more strongly, while higher values recovered higher numbers of ESVs. The default, and the value used in most studies with ribosomal DNA, is 2. However, for COI two independent approaches, based on mock communities (Elbrecht et al 2018) and entropy changes (Turon et al 2020) suggested that for this marker α=5 is the optimal value. For DADA2 the key parameter is omega_A, which indicates the probability than a sequence *i* is an error derived from another sequence *j* given their abundance values and the inferred error rates. If the observed value is higher than omega_A, then sequence *i* is considered an error of sequence *j*. Omega_A is by default set to a very low value (10^−40^), but no study has analysed the impact of changing this parameter for COI datasets. To our knowledge, only Tsuji et al (2020), based on a comparison of 3 values, concluded that the default value of Omega_A was adequate for a marker based on the control region of the mitochondrial DNA.

### The clustering algorithm

Our preferred clustering method is SWARM v3 (Mahé et al 2015), as it is not based on a fixed distance threshold and is independent of input order. It is a very fast procedure that relies on a single-linkage method with a clustering distance (*d*), followed by a topological refining of the clusters using abundance structures to divide MOTUs. As we were interested in keeping all sequences within MOTUs, and not just a representative sequence, we mined the SWARM output with an R script to generate MOTU files, each with its sequence composition and abundance.

The crucial parameter in this approach is *d*, the clustering distance threshold for the initial phase. The default value is 1 (that is, amplicons separated by more than one difference will not be clustered together), and this value has been tested in microbial ribosomal DNA. However, Mahé et al (2014) pointed out that higher *d* values can be necessary for fast evolving markers (such as COI) and advised to analyse a range of *d* to identify the best fitting parameter for a particular dataset or scientific question. A *d* value of 13 (thus, allowing 13 differences over ca. 313 bp to make a connection) has been recently used for the Leray fragment of COI (e.g. Siegenthaler et al 2019, Garcés-Pastor et al 2019, Bakker et al 2019, Atienza et al 2020, Antich et al 2020), but a formal study of its adequacy has not been published yet.

### Setting the right parameters

With our dataset, we assessed the best-fitting parameters for UNOISE3, DADA2 and SWARM as applied to COI data. For the first two, we used changes in diversity values per codon position (measured as entropy, Schmidt & Herzel 1997), as calculated with the R package *entropy* (Hausser & Strimmer 2009). Coding sequences have properties that can be used in denoising procedures (Turon et al 2020, Tsuji et al 2020). They have naturally a high amount of variation concentrated in the third position of the codons, while errors at any step of the metabarcoding pipeline would be randomly distributed across codon positions. Thus, examining the change in entropy values according to codon position can guide the choice of the best cleaning parameters. Turon et al (2020) suggested to use the entropy ratio (Er) between position 2 of the codons (least variable) and position 3 (most variable). In a simulation study these authors showed that Er decreased as more stringent denoising was applied until reaching a plateau, which was taken as the indication that the right parameter value had been reached.

Using the Er to set cut-points, we re-assessed the adequate value of α in UNOISE3 testing the interval of α=1 to 10. With the same procedure, we tested DADA2 for values of Omega_A between 10^−0.05^ (ca. 0.9) and 10^−90^.

For SWARM, we compared the output of SWARM with a range of values of *d* from 1 to 30 applied to our dataset (prior to denoising). We monitored the number of MOTUs generated and the mean intra- and inter-MOTU distances to find the best-performing value of *d* for our fragment.

### The impact of the steps and their order

With the selected optimal parameters for each method, we combined the two denoising procedures and the clustering step in different orders. We therefore combined denoising (Du for UNOISE3 algorithm implemented in DnoisE, Da for DADA2) and clustering with SWARM (S) and generated and compared datasets of ESVs and MOTUs as follows (for instance, DaS means that the dataset was first denoised with DADA2, then clustered with SWARM):

ESVs: Du, Da

MOTUs: DuS, DaS, SDu, SDa

For comparison of datasets, we used Venn diagrams and an average match index of the form

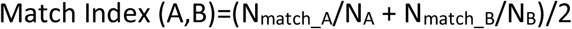

Where N_match_A_ is the number of a particular attribute in dataset A that is shared with dataset B, and N_A_ is the total number of that attribute in dataset A. The same for N_match_B_ and N_B_. The matches can be the number of ESVs shared, the number of MOTUs shared, the number of ESVs in the shared MOTUs, or the number of reads in the shared ESVs or MOTUs, depending on the comparison.

### Improving the denoising algorithm

The preferred denoising algorithm (UNOISE3, see Results) has been further modified in two ways. This procedure is based on two parameters: the number of sequence differences (d, as measured by the Levenshtein distance) and the abundance skew between two sequences (β), formalized in the simple formula (Edgar 2016):

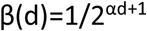

Where β(d) is the maximum abundance skew allowed between two sequences separated by distance d so that the less abundant was merged with the more abundant. Presumably incorrect daughter sequences are merged with the correct mother sequences if the number of sequence differences (d) is small and the abundance of the incorrect sequence with respect to the correct one (abundance skew) is low. The higher the number of differences, the lower the skew should be for the sequences to be merged.

The present algorithm of UNOISE3 gives precedence to the abundance skew over the number of differences (d) because sequences are considered in order of decreasing abundance. Thus, a very abundant sequence will form a centroid that can “capture” a rare one even if d is relatively high. Other, somewhat less abundant, sequences can be more similar (less d) to the rare sequence and can fulfil the conditions to capture it, but this will never happen as the rare sequence will be incorporated to the first centroid and will become unavailable for further comparisons. In our modification, DnoisE does not automatically join sequences to the first centroid that fits the condition. Rather, for each sequence the potential “mothers” are stored (with their abundance skew and d) and the sequences are left in the dataset. After the round of comparisons is completed, for each daughter sequence we can choose, among the potential mothers, the one whose abundance skew is lower (precedence to abundance skew, corresponding to the usual UNOISE3 procedure), the mother with the lowest distance (precedence to d), or the one for which the ratio (abundance skew/max abundance skew for the observed d, β(d)) is lower, thus combining the two criteria.

Second, for COI the fact that it is a coding gene is a fundamental difference with respect to ribosomal genes. In a coding fragment, the amount of variability is substantially different among codon positions. This is not taken into account in the UNOISE3 formulation (nor in DADA2 or other denoising programs that we knew of, for that matter). We suggest to incorporate this information in DnoisE by differentially weighting the d values according to whether the change occurs in the first, second, or third codon position. Note that our sequences are all aligned and without indels, which makes this weighting scheme straightforward. The differences in variability can be quantified as differences in entropy values (Schmidt and Herzel 1997); position 3 of the codons has the highest entropy, followed by position 1 and position 2. In other words, two sequences separated by n differences in third positions are more likely to be naturally-occurring sequences than if the n differences happen to occur in second positions, because position 3 is naturally more variable. To weight the value of d, we first record the number of differences in each of the three codon positions d(1) to d(3), we then change the d values using the formula

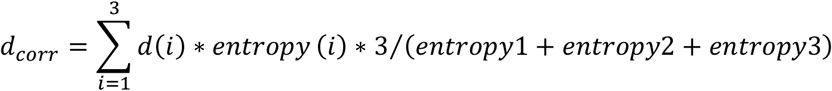

where d_corr_ is the corrected distance and *i* is the position in the codon.

With this formula, two sequences separated by just one difference in each codon position will continue to have a d of 3, but a change in a high entropy position (3) will translate in a higher d than the same change in a low entropy position (2), thus the program will tend to keep the former and to merge the later. The entropy of the three positions of the codons for the weighting scheme is taken from the study of Turon et al (2020) on 1,000 presumably correct COI sequences from a similar dataset of benthic communities, thus entropy(1)=0.430 bits, entropy(2)=0.183 bits, and entropy(3)=0.926 bits. Note that d_corr_ is based on the number of differences occurring at each codon position. The Levenshtein distance used in the non-corrected d measures is not adequate for this purpose, as it cannot keep track of codon positions. However, for sequences of equal length, aligned, and without indels, as in our case, the number of differences is practically equivalent to the Levenshtein distance.

We compared in our dataset the results of the different formulations of DnoisE: precedence to abundance skew, precedence to distance, combined precedence, and correcting distances according to codon position of the differences.

A beta version of DnoisE (<u>D</u>istance de<u>nois</u>e by <u>E</u>ntropy) is available from https://github.com/adriantich/DnoisE.

### Taxonomic validation

Ground truthing is a difficult task in metabarcoding studies. Constructing mock communities is the most common method. However, mock communities, even the largest ones, are orders of magnitude simpler than complex biological communities. Thus, some technical aspects cannot be tested accurately. For instance, metabarcoding results of mock communities in general lack true sequences at very rare abundances (the most problematic ones). We devised a checking procedure by taxonomically assigning our ESVs using the ecotag procedure in obitools against the db-COI_MBPK database (Wangensteen et al 2018), containing 188,929 eukaryote COI reference sequences (available at https://github.com/metabarpark/Reference-databases). Ecotag assigns a sequence to the common ancestor of the candidate sequences selected in the database, using the NCBI taxonomy tree. This results in differing taxonomic rank of the assignments depending on the density of the reference database for a given taxonomic group. As MOTUs should ideally reflect species-level entities, we selected those sequences assigned at the species level as a benchmark for the MOTU datasets. We traced these sequences in the output files of our procedures and classified the MOTUs containing them into three categories (following the terminology of Forster et al 2019): closed MOTUs, when they contain all sequences assigned to a species and only those; open MOTUs, when they contain some, but not all, sequences assigned to one species and none from other species, and hybrid MOTUs, when they contain sequences assigned to more than one species.

This analysis was intended as a tool for comparative purposes, to benchmark the ability of the different MOTU sets generated to recover species-level entities.. Our rationale is that a procedure that maximizes the relative number of closed MOTUs and minimizes hybrid MOTUs reflects more accurately the real biological variation in the samples.

## RESULTS

### The dataset

After pairing, quality filters, and retaining only 313 bp-long reads, we had a dataset of 16,325,751 reads that were dereplicated into 3,507,560 unique sequences. After deleting singletons (sequences with one read), we kept 423,164 sequences (totalling 10,305,911 reads). Of these sequences, 92,630 were identified as chimeras and 152 as misaligned sequences and eliminated. Out final dataset for the study, therefore, comprised 330,382 sequences and 9,718,827 reads (deposited in Mendeley Data, https://data.mendeley.com/datasets/84zypvmn2b/2).

For testing the performance of DADA2 on unpaired and paired reads on a coherent dataset, we selected the reads that were in the forward direction, that is, the forward primer was in the forward read (R1). As expected, they comprised half of the reads (4,892,084). For these reads we compared the output of applying DADA2 before and after merging, as detailed in Supplementary Material S1. The results were highly similar, with most reads placed in the same ESVs in both datasets, albeit 21% more low-abundance ESVs were retained using the paired reads. Henceforth we will use DADA2 on paired sequences, as results were markedly similar and this was necessary to perform our comparisons.

### Setting the right parameters

We used the change in entropy ratio (Er) of the retained sequences of the global dataset (330,382 sequences and 9,718,827 reads) for selecting the best performing α– value in UNOISE3 and the best Omega_A in DADA2 across a range of values. We also assessed the number of ESVs resulting from the procedures.

For UNOISE3 as implemented in our DnoisE script, the Er diminished sharply for α– values of 10 to 7, and more smoothly afterwards (Fig. 1A). The number of ESVs detected likewise decreased sharply as with lower α–values, but tended to level off at α=5 (Fig. 1A). The value of 5 seems a good compromise between minimizing the Er and keeping the maximum number of putatively correct sequences.

**Fig. 1.**
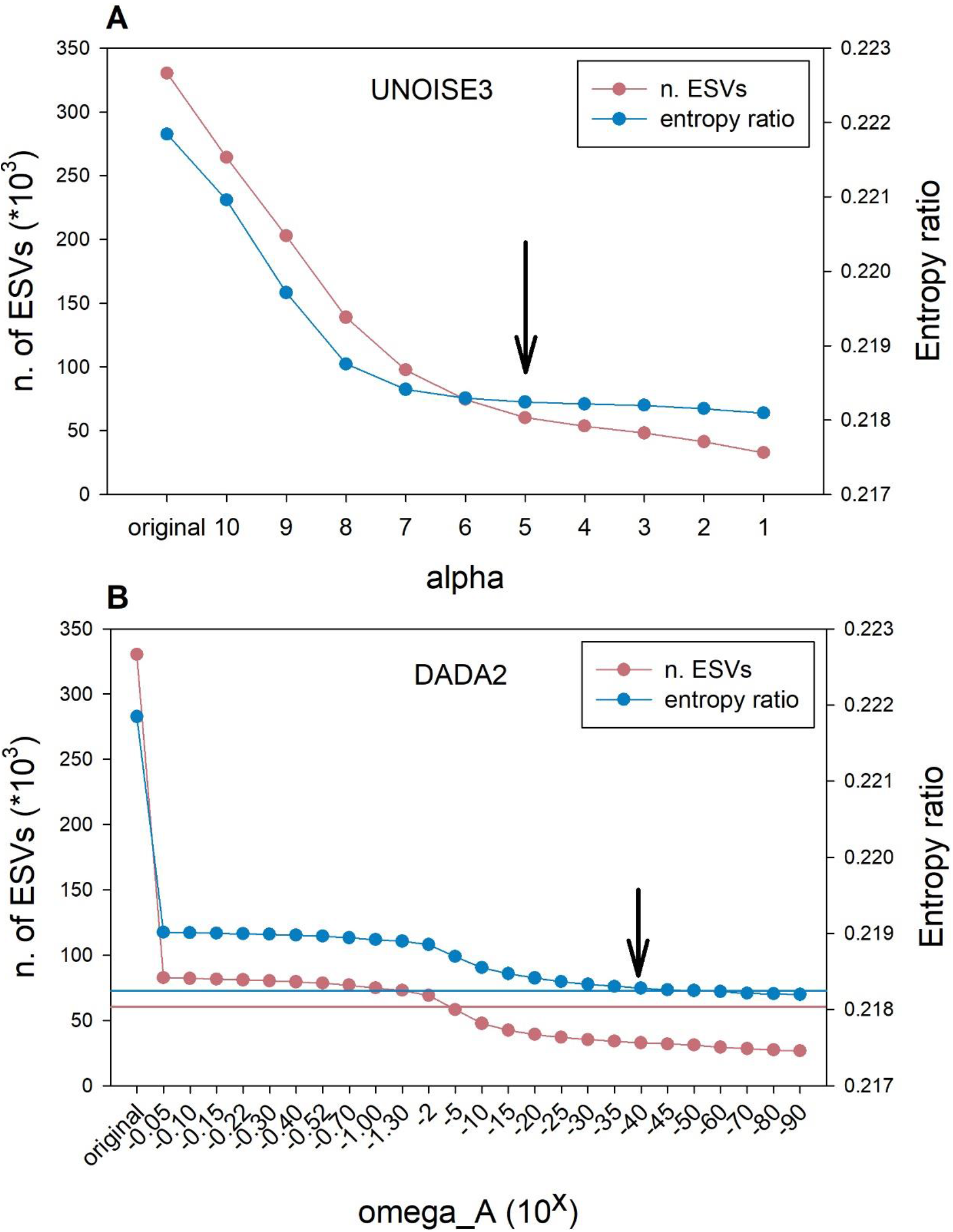
Values of the Entropy ratio (Er) of the set of ESVs obtained with the UNOISE3 algorithm at decreasing values of α (A), and of those obtained with the DADA2 algorithm at decreasing values of omega_A (B). Arrows point at the selected value for each parameter. Horizontal blue line in (B) represents the Er value reached in A at α=5, horizontal red line marks the number of ESVs detected in A at α=5.

For the DADA2 algorithm we tested a wide range of omega_A from 10^−0.05^ to 10^−90^ (we set parameter omega_C to 0 in all tests, so all erroneous sequences were corrected). The results showed that, even at the highest value (10^−0.05^, or ca. 0.9 p-value, thus accepting as new partitions an exaggerate number of sequences), there was a substantial drop in number of sequences (ca. 75% reduction) and in Er with respect to the original dataset (Fig. 1B). Both variables remained relatively flat with a slight decrease between omega_A 10^−2^ and 10^−15^, becoming stable again afterwards (Fig 1B).

The number of ESVs retained was considerably lower than for UNOISE3. In fact, the number obtained at α=5 by the latter (60,209 ESVs) was approximately reached at omega_A=10^−5^ (58,191 ESVs). On the other hand, the entropy value obtained at α=5 in UNOISE3 (0.2182) was not reached until omega_A=10^−60^. As a compromise, we will use in this study the default value of the dada function (10^−40^), while acknowledging that the behaviour of DADA2 with changes in omega_A for the parameters analysed was unexpected and deserves further research.

For the clustering algorithm SWARM v.2, we monitored the outcome of changing the *d* parameter between 1 and 30. For each value, we tracked the number of clusters formed (separately for all MOTUs and for those with 2 or more sequences), as well as the mean intra-MOTU and the mean inter-MOTU distances (considering only the most abundant sequence per MOTU). The goal was to find the value that maximizes the intra-MOTU variability while keeping a sharp difference between both values (equivalent to the barcode gap).

The total number of MOTUs decreased sharply from 38,560 (*d*=1) to around 1,900 with a plateau from *d*=9 to *d*=13, and then decreased again (Fig. 2A). If we only consider the MOTUs with 2 or more sequences, the overall pattern is similar, albeit the curve is much less steep. The numbers decreased from 8,684 for *d*=1 to 6,755 at *d*=12 and 13, and decreasing again at higher values (Fig. 2A).

**Figure 2.**
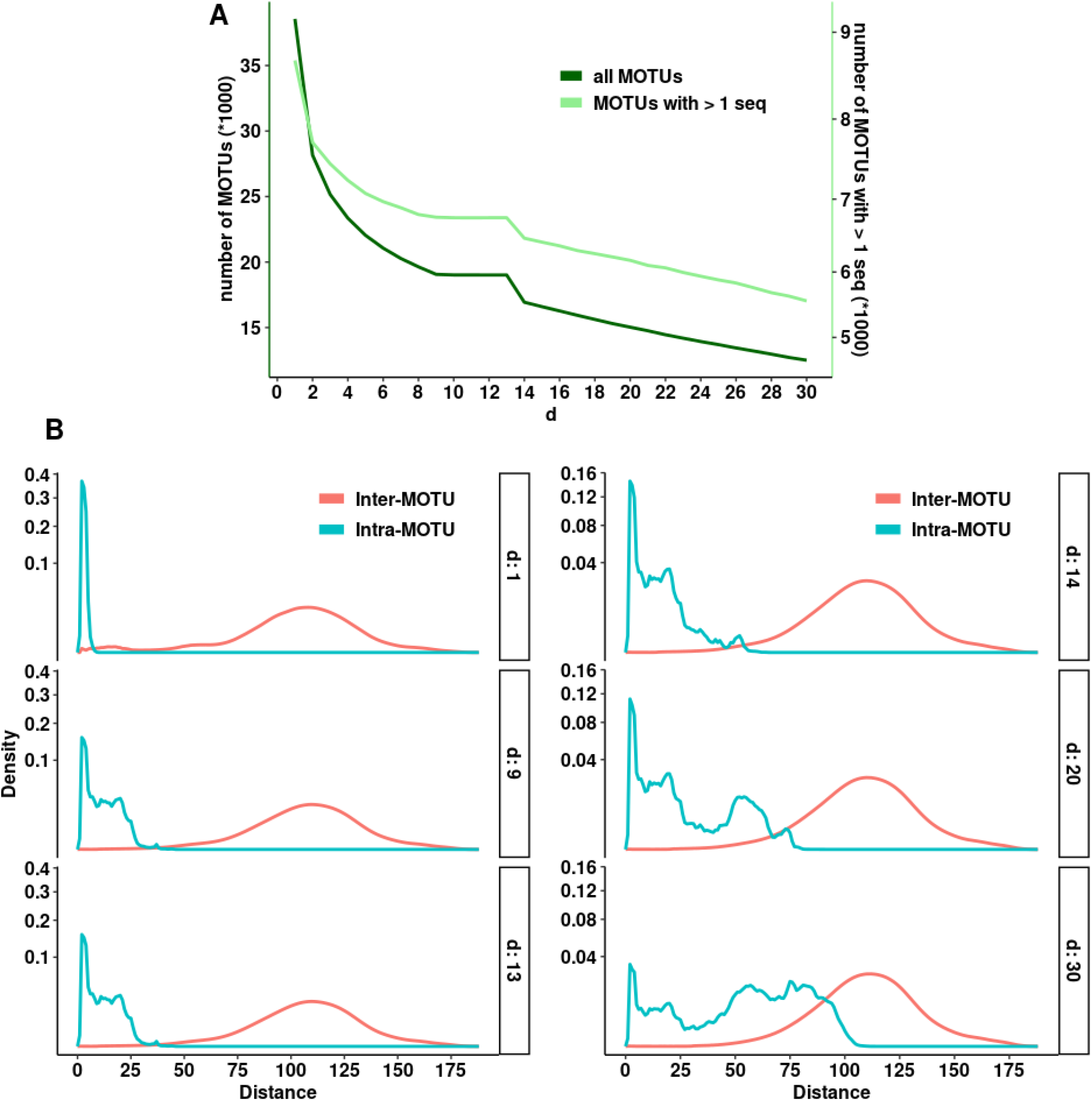
A. Number of MOTUs obtained at different values of d using SWARM. Total number of MOTUs (dark green) and of MOTUs with two or more sequences (light green) are represented (note different Y-axes). B. Density plots (note quadratic scale) showing the distribution of number of differences between different clusters (inter-MOTU, red) and sequences within clusters (intra-MOTU, blue) obtained by SWARM for selected values of the parameter d (1, 9, 13, 14, 20 and 30).

Inter-MOTU distances had a similar distribution with all values of the parameter *d*, albeit with a small shoulder at distances of 10-20 differences with *d*=1 (selected examples in Fig. 2B). Intra-MOTU distances, on the other hand, became more spread with higher values of *d* as expected. Values from 9 to 13 showed a similar distribution of number of differences, but for *d* values higher than 14, intra-MOTU distances started to overlap with the inter-MOTU distribution (Fig. 2B). The value of *d*=13 seems, therefore, to be the best choice to avoid losing too much MOTU variability (both in terms of number of MOTUs and intra-MOTU variation), and at the same time keeping intra- and inter-MOTU distances well separated. The mean intra-MOTU distance in our dataset at d=13 was 9.10 (equivalent to 97.09% identity), and the mean inter-MOTU distance was 108.78 (65.25% identity).

### The impact of the steps and their order

Table 1 shows the main characteristics of the original and the generated datasets, as well as the datasets obtained by modifying the UNOISE3 algorithm (see below). All datasets are available from Mendeley Data (https://data.mendeley.com/datasets/84zypvmn2b/2).

**Table 1.** Main characteristics of the original and the generated datasets. All datasets had 9,718,827 reads. 1-ESV MOTUs refer to the number of MOTUs with just one ESV. (*) for the original and S datasets the number of sequences instead of ESVs is used.

|  | n. ESVs * | n. MOTUs | 1-ESV<br>MOTUs | ESVs/MOTU * | reads/MOTU |
| --- | --- | --- | --- | --- | --- |
| Original | 330,382 | -- | -- | -- | -- |
| Du | 60,198 | -- | -- | -- | -- |
| Da | 32,798 | -- | -- | -- | -- |
| S | 330,382 | 19,012 | 12,257 | 17.378 | 511.194 |
| DuS | 60,198 | 19,058 | 12,471 | 3.159 | 509.961 |
| SDu | 75,069 | 19,012 | 12,433 | 3.949 | 511.194 |
| DaS | 32,798 | 19,167 | 15,565 | 1.711 | 507.060 |
| SDa | 35,376 | 19,012 | 15,198 | 1.861 | 511.194 |
| DdS | 60,198 | 19,058 | 12,471 | 3.159 | 509.960 |
| DcS | 60,198 | 19,058 | 12,471 | 3.159 | 509.960 |
| DeS | 82,856 | 19,089 | 12,609 | 4.341 | 509.132 |
| DedS | 82,856 | 19,089 | 12,609 | 4.341 | 509.132 |
| DecS | 82,856 | 19,088 | 12,608 | 4.341 | 509.159 |

We first compared the outcomes of denoising the original reads with UNOISE3 and DADA2 (Du vs Da), with the stringency parameters set as above. The error rates of the different substitution types as a function of quality scores were highly correlated in the DADA2 learnErrors procedure. The lowest Pearson correlation was obtained between the substitutions T to C and A to G (*r*=0.810), and all correlations (66 pairs of substitution types) were significant after a False Discovery Rate correction (Benjamini & Hochberg 1995).

The main difference found is that the Du dataset retained almost double number of ESVs than the Da dataset: 60,198 vs 32,798. Of these, 31,696 were identical in the two datasets (Fig. 3), representing a match index of 0.746. Of the shared ESVs, 20,691 (65.28%) had exactly the same number of reads, suggesting that the same reads have been merged in these ESVs.

**Figure. 3.**
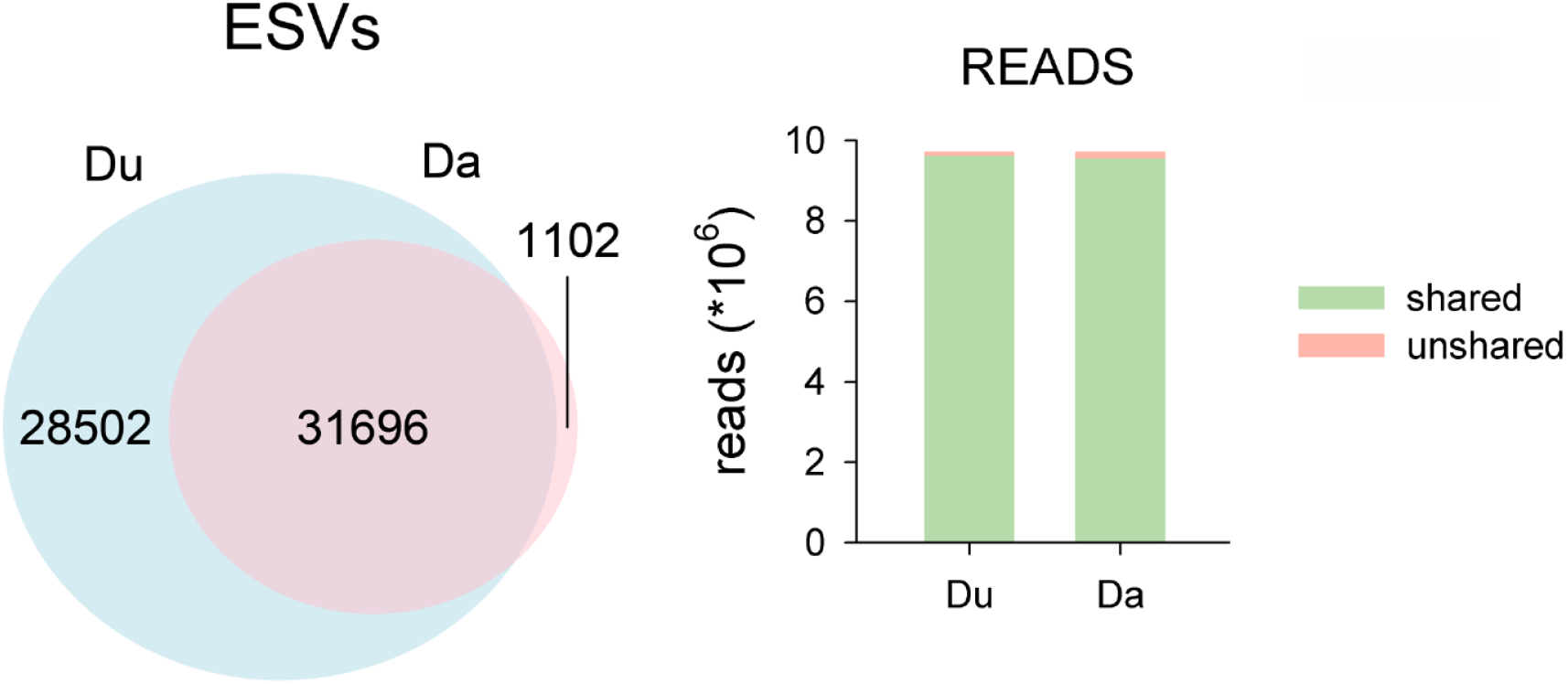
Venn Diagram showing the number of ESVs shared between the two denoising procedures before (Du vs Da). Bar chart shows the number of reads in the shared and unshared ESVs.

On the other hand, the shared ESVs concentrated most of the reads (Fig. 3): the match index for the reads was 0.986. This is coherent with the fact that most of the non-shared ESVs of the Du dataset had a low number of reads (mean=3.66).

Thus, the two denoising algorithms with the chosen parameter values provided similar results as for the abundant ESVs, but UNOISE3 retained a high number of low abundance ESVs as true sequences.

We then evaluated the output of combining denoising and clustering, using either of them as a first step. Thus, we compared the datasets DuS, SDu, DaS, and SDa.

The results showed that the final number of MOTUs obtained was similar (ca. 19,000) irrespective of the denoising method and the order used (Table 1). Moreover, the shared MOTUs (flagged as MOTUs that have the same representative sequence) were the overwhelming majority (Fig. 4), with MOTU match indices over 0.96 in all comparisons.

**Figure. 4.**
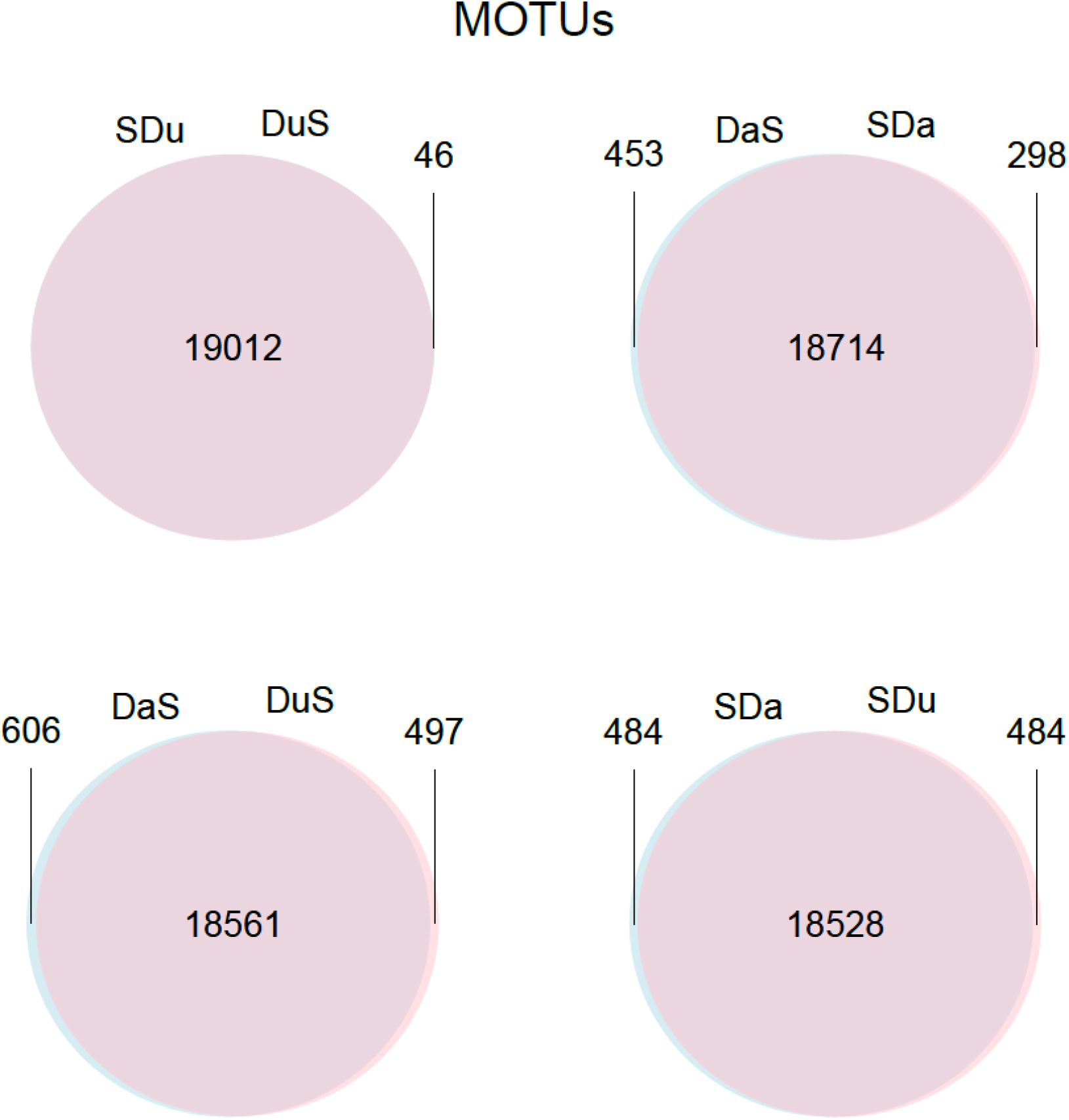
Venn Diagrams showing the number of MOTUs shared between the two denoising procedures and a clustering step performed in different orders.

As for the number of ESVs, clustering first results in a higher number of retained sequence variants than clustering last, ca. 25% more for Du and ca. 8% for Da. In all comparisons, the majority of ESVs were to be found in the shared MOTUs, and the same applies to the number of reads (Fig. 5, match indices for the ESVs, all >0.95, match indices for the reads, all > 0.99). Ca. 2/3 of the MOTUs comprised a single ESV when using Du, and this number increased notably with Da (ca. 80% of MOTUs, Table 1). In both cases, clustering first resulted in a slight decrease of the number of single-ESV MOTUs

**Figure. 5.**
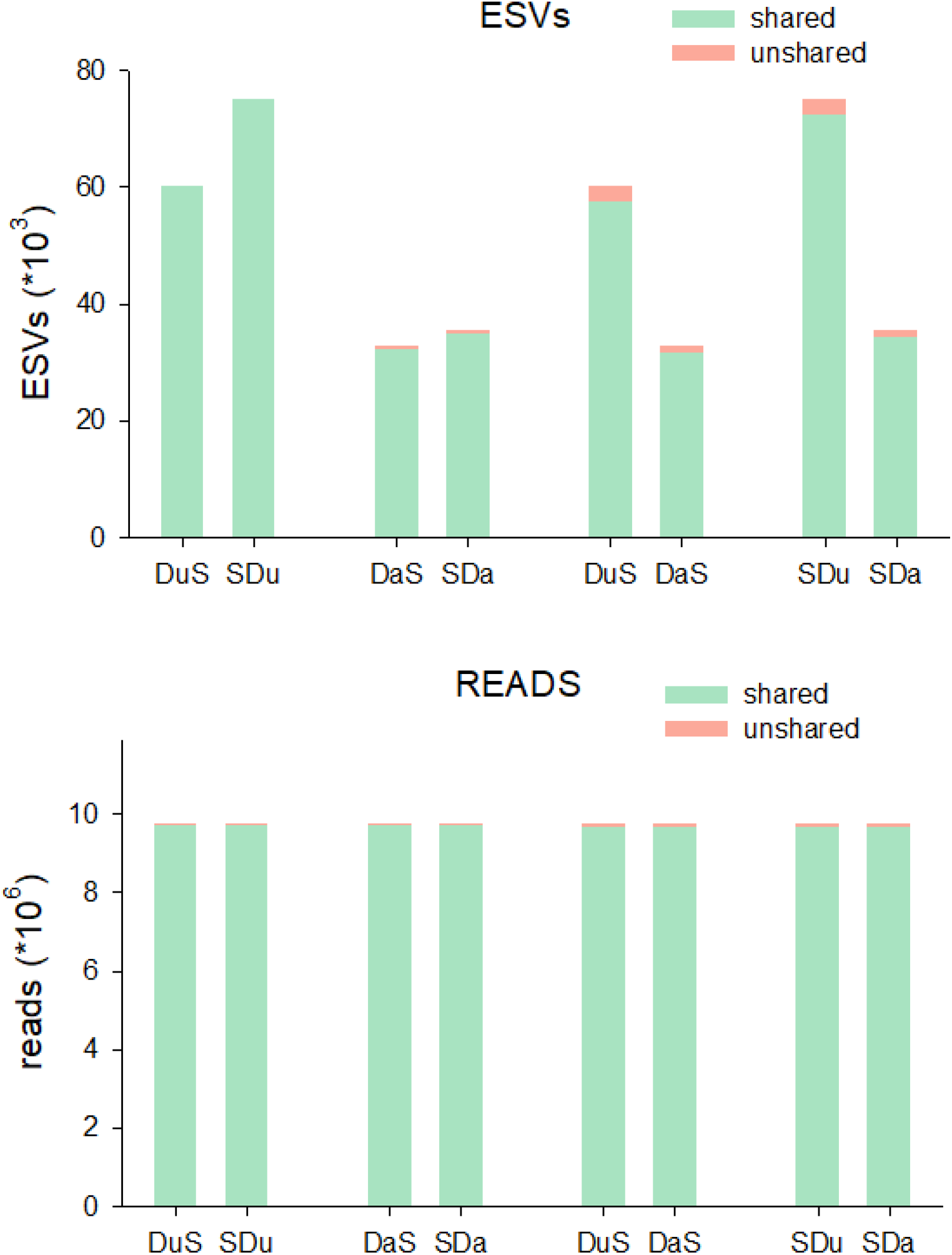
Bar charts of the number of ESVs and reads in the MOTUs shared or unshared in the same comparisons as in Figure 4.

### Improving the denoising algorithm

We tried different options of our DnoisE algorithm. The use of the Levenshtein distance without any correction and with priority to abundance skew corresponds to the original UNOISE3 algorithm (i.e., the Du dataset used previously). We also tried priority to <u>d</u>istance (Dd) and a <u>c</u>ombination of skew and abundance (Dc) to decide among the potential “mother” ESVs to which a given “daughter” sequence will be joined. The same three options were applied when correcting distances according to the <u>e</u>ntropy of each codon position (datasets De, Ded, and Dec). In this case we used a pairwise distance accounting for the codon position where a substitution was found. We further applied a clustering step (SWARM) to the DnoisE results to generate MOTU sets (DuS, DdS, DcS, DeS, DedS, DecS) for comparison with those obtained previously.

The three ways to join sequences have necessarily the same ESVs, only the sequences that are joined under each centroid can vary and, thus, the abundance of each ESV. However, this had a very small effect. For the three datasets generated without distance correction (the original Du dataset, plus Dd and Dc), over 99% of ESVs have the same number of reads, suggesting that the same sequences have been grouped in each ESV. For the entropy-corrected datasets (De, Ded, Dec), the number is slightly lower (over 96%).

On the other hand, if we take into account the entropy of codon positions the results change notably. The corrected datasets have 82,856 ESVs (against 60,209 of the uncorrected datasets). So, when considering the entropy in distance calculations the number of retained ESVs increased by ca. 38%. This is the result of accepting sequences that have variation in third codon positions as legitimate. Further, the sequences retained were also different: when comparing the entropy-corrected and uncorrected datasets only 46,626 ESVs were found in common (ESV match index of 0.669). These ESVs comprise a majority of reads, though (read match indices of ca. 0.97 in all possible comparisons). Fig. 6 illustrates one of these comparisons (Du vs Dec).

**Figure 6.**
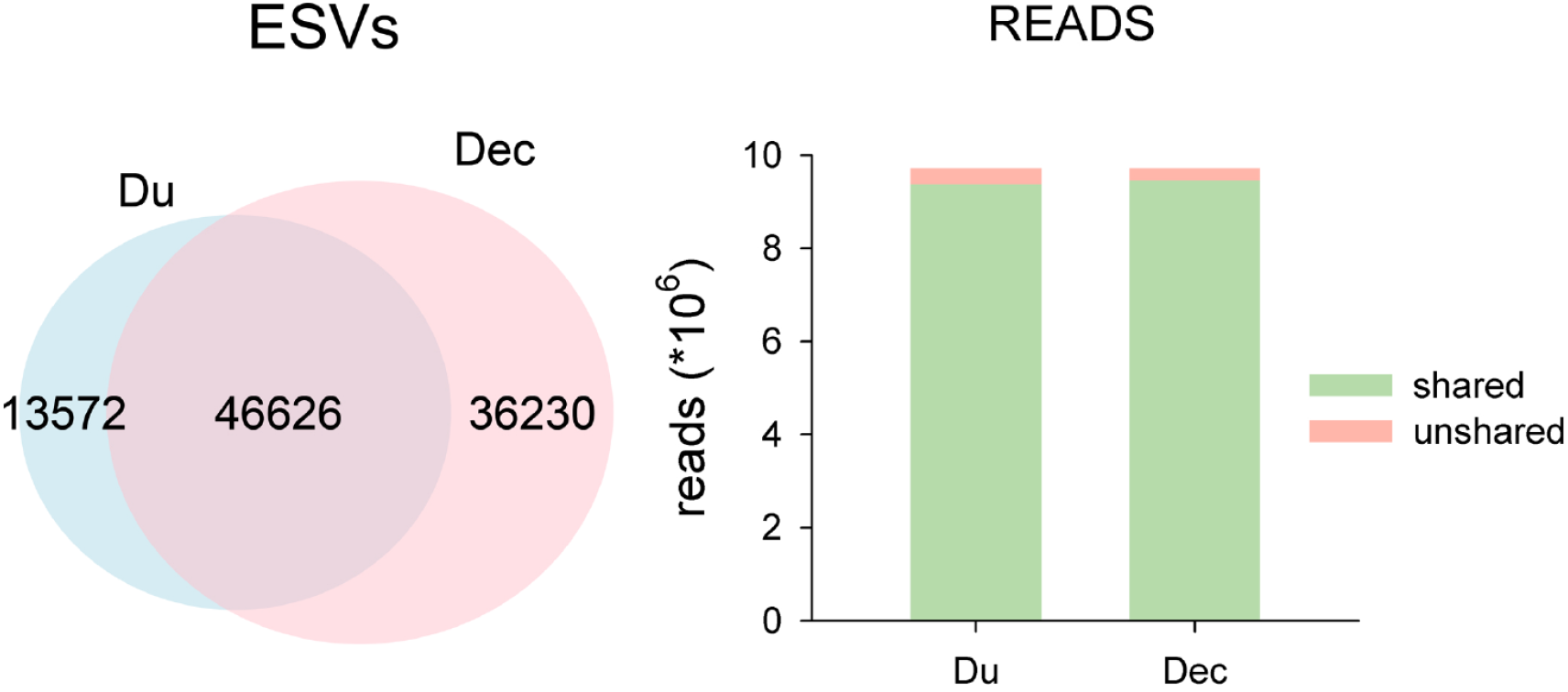
Venn Diagram showing the number of ESVs shared between two denoised datasets (Du vs Dec). Bar chart shows the number of reads in the shared and unshared ESVs.

When clustering the ESVs obtained with the different methods, the final number of MOTUs obtained was similar to those generated in the previous sections (ca. 19,000 in all cases, Table 1). This indicates that the entropy corrected datasets provided more intra-MOTU variability, but no appreciable increase in the number of MOTUs. As an instance, the mean number of ESVs per MOTU was 3.159 for the DuS dataset, and 4.341 for the DecS dataset. Only a slight increase in the number of single-ESV MOTUs was detected (12,471 for DuS, 12,608 for DecS). We show in Figure 7 the comparison between these two datasets. Most MOTUs (as indicated by identity in the representative sequence) were shared between datasets. In addition, most of the ESVs and most of the reads were found in the shared MOTUs (match ratios for MOTUs, ESVs and reads >0.99. Fig. 7).

**Figure 7.**
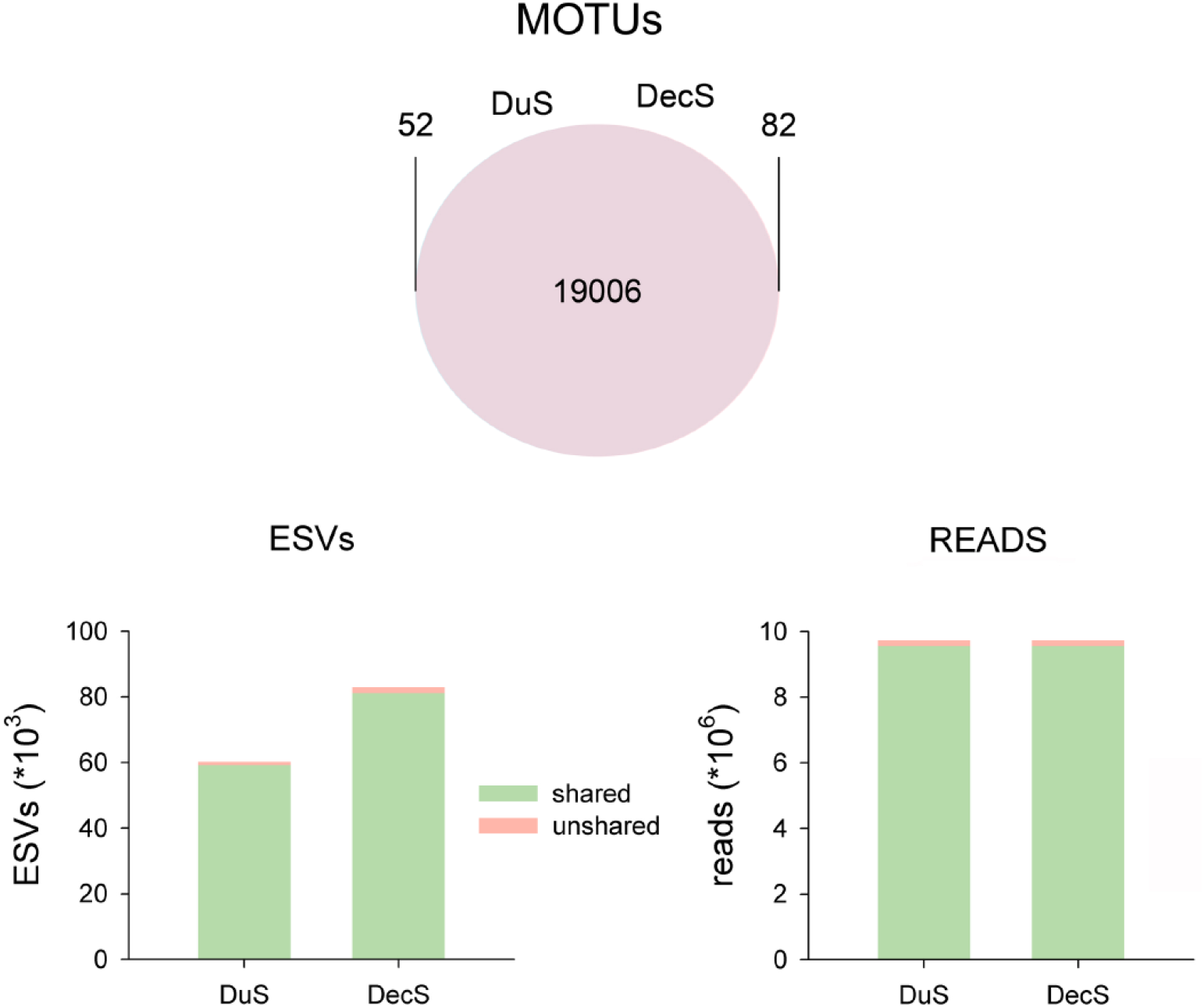
Venn Diagrams showing the number of MOTUs shared between datasets DuS and DecS. Bar charts show the number of ESVs and reads in the shared and unshared MOTUs.

### Taxonomic validation

We combined all unique ESVs retained after the denoising algorithms (those retained by the different versions of DnoisE and those retained by DADA2, for a total of 97,379 ESVs) and assigned them taxonomically with ecotag. We found that 19,108 sequences had a species-level assignment, comprising 681 species, of which 189 were represented by a single sequence. We further refined this dataset by accepting only sequences whose best hit in the reference database was ≥ 0.95. This pruned dataset (henceforth species-level dataset, available as Table S1), consisted of 12,251 assigned sequences and 447 species, with 142 having only one sequence. The inclusion of the entropy-corrected ESV datasets (which kept more ESVs than the other methods) doubled the number of sequences with species-level hit but these represented only one extra species with respect to the other datasets, again indicating that the gain in ESVs in the entropy-corrected procedure mainly increases within-MOTU variability.

We checked if the sequences in the species-level dataset were grouped in closed MOTUs (meaning all sequences in the MOTU belonged to the same species and no other sequences of this species were found in other MOTUs), open MOTUs (i.e., all sequences belonged to the same species, but not all sequences of the species were included) and hybrid MOTUs (i.e., including sequences assigned to more than one species, or a combination of sequences assigned to species and sequences unassigned at this rank). Closed MOTUs were further subdivided among those with only one sequence (closed singleton) and those with several sequences (closed group).

As shown in Figure 8, when we checked the different datasets generated with the two denoising algorithms and different processing steps, irrespective of the method ca. 57% of MOTUs that had sequences assigned to the species rank were closed and ca. 25% open, with only ca. 18% hybrid MOTUs. This indicates that, in all cases, the denoising plus clustering methods performed well in recovering species that were identified as such in the ESVs. The UNOISE3 algorithm, however, recovered almost double closed group MOTUs that DADA2, and the opposite occurred for closed singleton MOTUs, for a similar total. This is the result of the higher number of sequences retained by UNOISE3, that translated into a higher ability to recover MOTUs with internal diversity. Differences were also apparent in the proportion of ESVs with species-level assignment that were found in the different datasets. In general, DADA2-based datasets had a lower proportion of ESVs with species assignment. Clustering first reduced appreciably (DuS vs. SDu, ca. 13% reduction) or marginally (DaS vs. SDa, 1.4%) this proportion. Entropy-corrected datasets had not only a higher number of ESVs, but a higher proportion of them (>11%) with species-level assignment.

**Figure 8.**
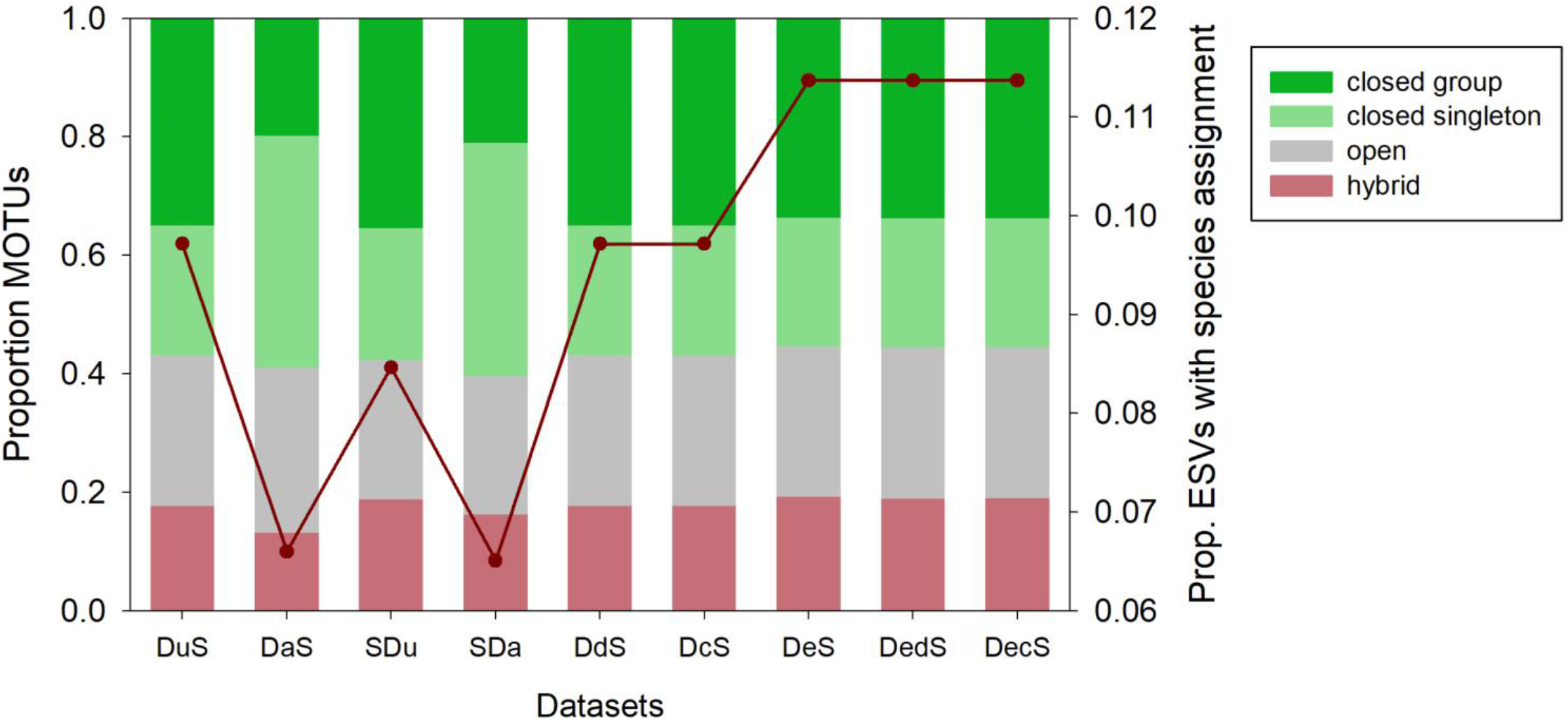
Proportion of MOTUs, among those that had any sequence identified at the species level, that were closed group, closed singleton, open, and hybrid. The red line shows the proportion of ESVs with species-level assignments found in each dataset.

## DISCUSSION

After adjusting the different parameters of the algorithms based on *ad hoc* criteria for COI amplicons, between ca. 33,000 and ca. 83,000 ESVs were obtained depending on the denoising procedure used. Irrespective of the method, however, they clustered into ca. 19,000 MOTUs. This implies that there was an important intra-MOTU variability even for the most stringent denoising method. The application of SWARM directly to the original dataset (without any denoising) generated likewise ca. 19,000 MOTUs. This suggests that the SWARM algorithm is robust in recovering alpha-diversity even in the presence of noisy sequences. Thus, denoising and clustering clearly accomplish different functions and, in our view, it is incorrect to use one instead of the other. The fact that some studies detect more MOTUs than ESVs when analysing datasets using clustering and denoising algorithms separately (e.g., Macheriotou et al 2018, Nearing et al 2028, Giebner et al 2020) reflects this logical flaw: MOTUs seek to recover meaningful species-level entities, ESVs seek to recover correct sequences. There should be more sequences than species, otherwise something is wrong with the respective procedures. It has even been suggested that ESVs or MOTUs represent a first level of sequence grouping and that a second round using network analysis is convenient (Forster et al 2019). We contend that, with the right parameter settings, this is unnecessary for eukaryotic COI datasets.

We do not endorse Callahan et al (2017) view that ESVs should replace MOTUs as the standard unit analysis of amplicon-sequencing datasets. Using information at the strain level may be useful in the case of prokaryotes, and in low-variability eukaryote markers such as ribosomal 18S rDNA there may be correspondence between species and unique sequences (indeed, in many cases different species share sequences). But even in more variable nuclear markers such as ITS, a clustering step is necessary (Estensmo et al 2020). In eukaryotes the unit of diversity analyses is the species, and in highly variable markers such as COI, it is well known that there is a rich intraspecies variability. MOTUs and not ESVs target species-level diversity and, in our view, should be used as the standard unit of analyses for most ecological and monitoring applications. Most importantly, that ESVs are organized into MOTUs is highly relevant information added at no cost. We do not agree that clustering ESVs into MOTUs eliminates biological information (Callahan et al 2016). This only happens if only one representative sequence per MOTU is kept. But it does not need to be so. We strongly advocate here for keeping track of the different sequences clustered in every MOTU and reporting them in metabarcoding studies. In this way analyses can be performed at the MOTU level or at the ESV level, depending on the question addressed.

Denoising has been suggested as a way to overcome problems of MOTU construction and to provide consistent biological entities (the correct sequences) that can be compared across studies (Callahan et al 2017). We fully agree with the last idea: ESVs are interchangeable units that allow comparisons between datasets and can avoid generating too big datasets when combining reads of, say, temporally repeated biomonitoring studies. But clustering ESVs into MOTUs comes as a bonus, provided the grouped sequences are kept and not collapsed under a representative sequence, thus being available for future reanalyses.

The denoising and clustering methods here tested have been developed for ribosomal markers and uncritically applied to COI data in the past, with default parameter values taken at face value (in fact, parameters are rarely mentioned in methods sections). We confirm that the UNOISE3 parameter α is best set at 5 for COI data, in agreement with Elbrecht et al (2018) and Turon et al (2020). We also confirm the suitability of a *d* value of 13 for SWARM that has been used in previous works with COI datasets (e.g., Siegenthaler et al 2019, Garcés-Pastor et al 2019, Bakker et al 2019, Atienza et al 2020, Antich et al 2020). As Mahé et al (2014) noted, high *d* values can be necessary for fast evolving markers. They advised to track MOTU coalescing events as *d* increases to find the value best-fitting the sequence marker chosen. We have used this information together with the intra- and inter-MOTU distances to select the *d*-value, which is clearly marker-dependent. In our view, fixed-threshold clustering procedures should be avoided, as even for a given marker the intra- and interspecies distances can vary according to the group of organisms considered. With SWARM, even if the initial clusters were made at d=13 (for a fragment of 313 this means an initial threshold of 4.15% for connecting sequences), after the refining procedure the mean intra-MOTU distances obtained was 2.91%, which is in line with values suggested using the whole barcoding region of COI (Ratnasingham & Hebert 2013). Furthermore, in our taxonomic checking, we found a low proportion of hybrid MOTUs, irrespective of the denoising method used, indicating that the SWARM procedure adequately and robustly grouped the sequences with known species-level assignments.

Our preferred algorithm for denoising is UNOISE3. It is a one-pass algorithm based on a greedy formula with few parameters, it is computationally fast and can be applied at different steps of the pipelines. It keeps almost double ESVs than DADA2 and, combined with a clustering step, results in less single-sequence MOTUs and a higher number of ESVs per MOTU, thus capturing a higher intra-MOTU diversity. It also produced twice as much closed group MOTUs than DADA2 in our taxonomic validation. Edgar et al (2016), by comparing both algorithms in mock and *in vivo* datasets, also found that UNOISE had comparable or better accuracy than DADA2. Similarly, Tsuji et al (2020) found that UNOISE3 retained less false haplotypes than DADA2 in samples from tank water containing fish DNA. We found that the entropy values of the sequences changed as expected when denoising becomes more stringent with UNOISE3, indicating that the algorithm performs well with coding sequences. We also suggest ways of improving this algorithm (see below).

DADA2, on the other hand, is being increasingly used in metabarcoding studies but its suitability for COI remains to be demonstrated. We had to use paired reads (contrary to recommendation) to be able to make meaningful comparisons, but our results indicate that with unpaired sequences the number of ESVs retained would have been even lower. The DADA2 algorithm, when tested with increasingly stringent parameters, did not progressively reduce the entropy ratio values that should reflect an adequate denoising of coding sequences. Further, the high correlation of error rates between all possible substitution types suggests that the algorithm may be over-parameterized, at least for COI, which comes at a computational cost. Comparisons based on known communities (as in Tsuji et al 2020) and using COI are needed to definitely settle the appropriateness of the two algorithms for metabarcoding with this marker.

In addition, PCR-free methods now popular in library preparation procedures complicate the use of DADA2 as there is no consistent direction (forward or reverse) of the reads. We acknowledge that our merged sequences still included a mixture of reads that were originally in one or another direction and, thus, with different error rates. However, the non-overlapped part is only ca. 100 bp at each end of the sequences, and these are in general good quality positions in both the forward and reverse reads.

Another choice to make is to decide what should come first, denoising or clustering. Both options have been adopted in previous studies (note that clustering first is not possible with DADA2 unless paired sequences are used). Turon et al (2020) advocated that denoising should be made within MOTUs, as they provide the natural “sequence environment” where errors occur and where they should be targeted by the cleaning procedure. This is of course true only as long as the clustering process is error-free (i.e., no sequence is placed in the wrong cluster). This is unlikely to be the case, but the error rate of the clustering process is unknown. We found that clustering first retained more ESVs, because sequences that would otherwise be merged with another from outside its MOTU were preserved. It also resulted in less single-ESV MOTUs, retaining more intra-MOTU variability. It can also be mentioned that denoising the original sequences took approximately 10 times more computing time than denoising within clusters, which can be an issue depending on the dataset and the available computer facilities. We acknowledge, however, that most MOTUs are shared and most ESVs and reads are in the shared MOTUs when comparing the two possible orderings, irrespective of denoising algorithm. The final decision may come more from the nature and goals of each study. For instance, a punctual research may go for clustering first and denoising within clusters to maximize the intra-MOTU variability obtained. A long-term research that implies multiple samplings over time that need to be combined together may use denoising first and then perform the clustering procedure at each reporting period with the ESVs obtained in the datasets collected so far pooled.

We acknowledge that there are other important steps at which errors can be reduced and that require key choices (f.i., chimera identification, singleton treatment, unequal length sequences), but they are outside the scope of this work as we addressed only clustering and denoising steps. In particular, singletons (sequences with only one read) are a problem for all denoising algorithms (as it is difficult to discern rare sequences from errors). Singletons are often eliminated right at the initial steps, as we did in this work. However, it has been suggested that clustering first does not require prior elimination of singletons, as most will be placed in the correct cluster, and subsequent elimination of MOTUs consisting of a single read will alleviate the problem of excess singletons without eliminating all of them (Wangensteen et al 2018, Atienza et al 2020). Singletons remain an issue whose clarification requires further work. Likewise, a filtering step, in which ESVs with less than a certain amount of reads are eliminated, is deemed necessary to obtain biologically reliable datasets (Turon et al. 2020). However, this procedure and the adequate threshold are best adjusted according to the marker and the study system, so, albeit we acknowledge that a filtering step is likely necessary, this has not been addressed in this paper.

We recommend that the different denoising programs be programmed as stand-alone steps (not combined, for instance, with chimera filtering) so anyone interested could combine the denoising step with the preferred choices for other steps. We also favour open source programs that could be customized if needed. For UNOISE3 algorithm we suggest that a combination between distance and skew ratio be considered to assign a read to the most likely centroid. For DADA2 algorithm, we advise to weight the gain of considering the two reads separately vs using paired sequences. The advantages of the latter involve a higher flexibility of the algorithm as it does not need to be performed right at the beginning of the pipeline. For both algorithms, we think it is important to consider the natural variation of the three positions of the codons of a coding sequence such as COI, which can allow a more meaningful computation of distances between sequences and error rates. This of course applies to other denoising algorithms not tested in the present study (e.g., deblur Amir et al 2017, AmpliCI Peng& Dorman 2020). Our DnoiSE program, based on the UNOISE3 algorithm, includes the option of incorporating codon information in the denoising procedure. With this option, we found ca 20,000 more ESVs than with the standard approach; more importantly, ca. 36,000 ESVs obtained in the position-corrected datasets were not found in the non-corrected datasets. Albeit the shared ESVs contained most of the reads, there were important differences in the distribution of the less abundant ESVs. In our taxonomic checking, a noticeably higher proportion of ESVs with species-level matches in the reference database were detected with the codon position-corrected method. We used a dataset of fixed sequence length and eliminated misaligned sequences. The correction for codon position would be more complicated in the presence of indels and dubious alignments. We hope this approach will be explored further and adequately benchmarked in future studies.

In conclusion, we strongly advise, for variable markers such as COI, to combine denoising and clustering processes to make the most of the inter- and intraspecies information contained in metabarcoding datasets. We here provide recommendations for the preferred algorithms and step order, but these may be tuned according to the goals of each study, feasibility of preliminary tests, and ground-truthing options, if any. We feel, however, that it is highly recommendable to report the results in terms of both MOTUs and ESVs included in each MOTU, rather than reporting only MOTU tables with collapsed information and just a representative sequence. We also advise that the coding properties of COI should be used both to set the right parameters of the programs and to guide error estimation in denoising procedures. Keeping more ESVs may be a better option than being too stringent, because merging ESVs is always possible when more data or new evidence become available. Ultimately every researcher must decide the best option for their data. There is a huge amount of intra- and interMOTU information in metabarcoding datasets that can be exploited for basic (e.g., biodiversity assessment, connectivity estimates, metaphylogeography) and applied (e.g., management) issues in biomonitoring programs, provided the results are reported adequately.

## ACKNOWLEDGMENTS

We thank Daniel Sanromán for his help with the field sampling. The present research has been funded by Project PopCOmics (CTM2017-88080, MCIU/AEI/FEDER/UE) from the Spanish Government.

## AUTHOR CONTRIBUTIONS

XT and OSW conceived the study, XT, AA and CP performed the sampling, AA performed the laboratory work, AA and OSW wrote code, XT, CP, AA and OSW performed analysis, XT and AA drafted the ms, all authors contributed intelectual content and revised the ms.

## SUPPLEMENTARY FILES

Figure S1. Map of the sampling localities in the Iberian Peninsula, with indication of their coordinates. The map was generated with ggplot (Kahle D, Wickham H. ggmap: spatial visualization with ggplot2. The R Journal. 2013;5(1):144-161.

Supplementary Material S1. Comparison of DADA2 on paired and unpaired reads.

Table S1. S<u>pecies-level dataset.</u> Table with the ESVs identified at the species level with >95% similarity. The taxonomy assigned is indicatd, as well as the best-match in the reference database, the taxid information, and the sequence.

## Notes

https://data.mendeley.com/datasets/84zypvmn2b/2

https://github.com/adriantich/DnoisE

https://github.com/metabarpark/Reference-database

### Competing Interest Statement

The authors have declared no competing interest.

