## Supplementary figures and images for "To denoise or to cluster? That is not the question. Optimizing pipelines for COI metabarcoding and metaphylogeography"

### Figure S1

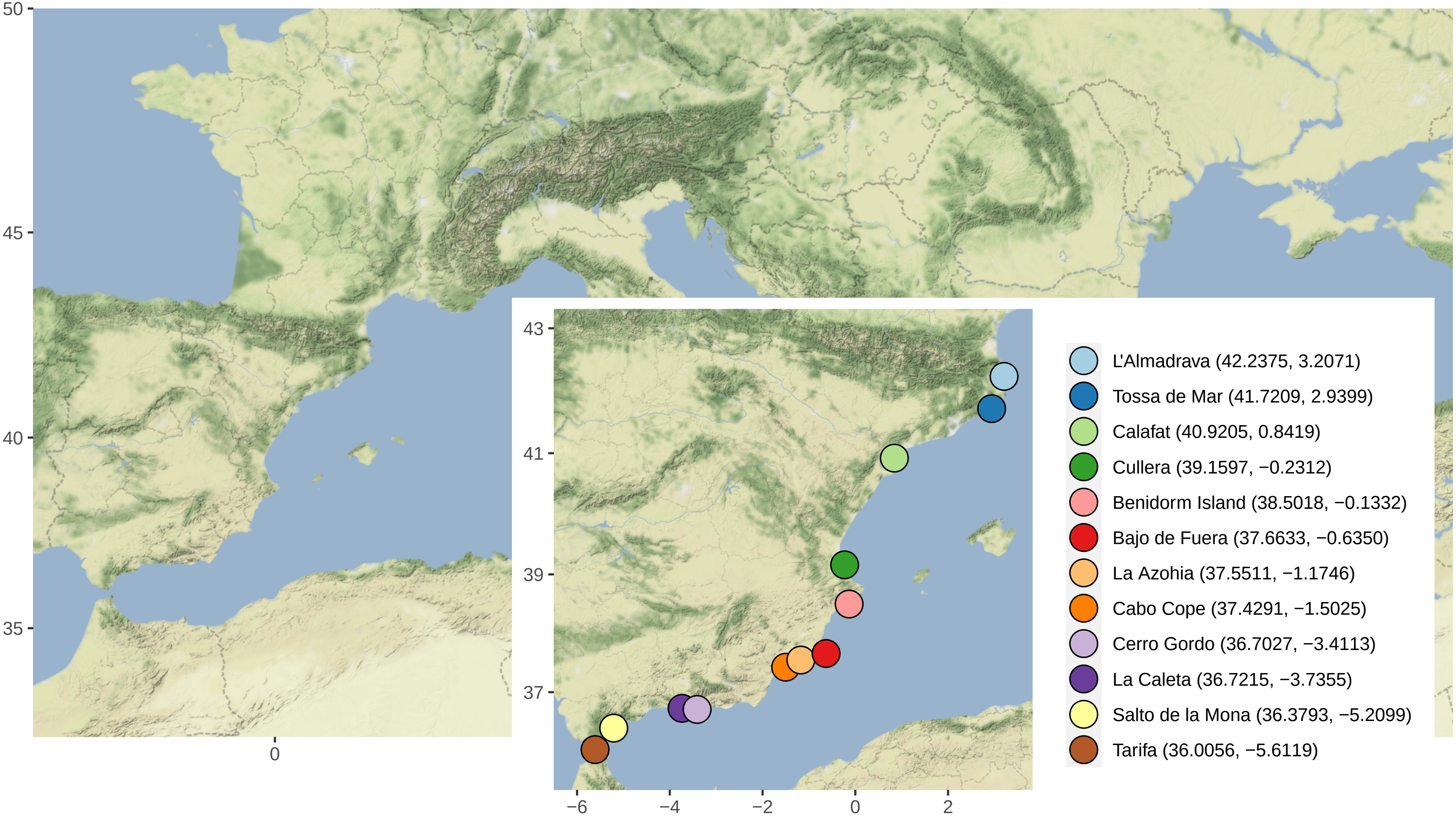
