## Supplementary material S1 for "To denoise or to cluster? That is not the question. Optimizing pipelines for COI metabarcoding and metaphylogeography"

**Comparison of DADA2 on unpaired and paired reads**

For testing the performance of DADA2 on unpaired and paired reads on a coherent dataset, we selected the reads that were in the forward direction, that is, the forward primer was in the forward read (R1). To do this, we selected the forward-oriented paired reads before de-replicating (as indicated by the tag "direction=forward" added by the merging procedure), but kept only those corresponding to sequences that will pass all the filters and will therefore make part of the final 9,718,827 reads.

As expected, the forward directed reads comprised ca. half of the total (4,892,084). This is due to the ligation-based library preparation protocol. We retrospectively picked the corresponding R1 and R2 reads from the sequencer output before pairing and eliminated the tags and primers. The last 20 bases from each read were trimmed. Thus, we had exactly the same 4,892,084 reads, paired and unpaired, for testing DADA2. This dataset, and the resulting ESV tables, are available from Mendeley Data (https://data.mendeley.com/datasets/84zypvmn2b/2).

We first applied DADA2 to the unpaired R1 and R2 reads using the R package dada2 v. 1.14, with error rates estimated from the data with learnErrors. The dada command was applied to R1 and R2 reads with the default value for omega_A ($10^{-40}$) and setting omega_C to 0 (so all sequences with errors were corrected) and DETECT_SINGLETONS to True (to use all reads). The resulting reads were merged with mergePairs. As a final output, we obtained 20,322 ESVs including the 4,892,084 reads.

The same procedure was repeated with the 4,892,084 paired reads. We input the sequences as if they were forward reads. The quality profiles showed the expected jump towards higher quality in the overlapped fragment (ca. 106 bp). The mean quality score of all positions is 51.29, of the non-overlapping positions is 39.66, and of the overlapping bases is 73.87. The error rates were computed from the data and dada was applied as before. The paired reads were input as forward reads, no reverse reads were input and no merging step was performed.

We obtained 24,573 ESVs, also totalling 4,892,084 reads. Therefore, using paired reads we obtained a 21% higher number of ESVs than with the unpaired reads. When comparing the outputs, we noted that 18,194 ESVs were identical (Fig 1). The match index of the ESVs was 0.818. In addition, the shared ESVs comprised most of the reads of the two datasets (98.81% of the reads of the unpaired dataset and 98.65% of the reads from the paired dataset). The match index of the reads was 0.987. The ESVs in the paired output not shared with the unpaired dataset had a low number of reads in general (average 10.39 reads).

We also noted that the estimated error rates reads for each substitution type (12 types) and quality score were highly correlated between the R1 and R2 reads ($r=0.870$, $p<10^{-15}$). In addition, the error rates as a function of quality score were also highly correlated between the 12 substitution types in each dataset. The lowest Pearson correlation coefficient for the estimated error rates of the R1 reads was 0.653 (between G to C and G to T changes), for R2 reads it was 0.741 (between G to T and C to G), and for the paired sequences it was 0.894 (between G to A and C to G). All correlations proved highly significant after a False Discovery Rate correction (Benjamini & Hochberg 1995).

Thus, our results using merged reads instead of using the forward and reverse sequences separately resulted in most reads being placed in the same ESVs, but more (21%) ESVs were kept when using merged reads. This result stems from the fact that a higher confidence in the bases of the long (ca. one third) overlapped region in turn results in accepting as correct sequences that otherwise would be labelled as erroneous. We could therefore retain low abundance ESVs that would otherwise be merged. Indeed, the ability to tell low-abundance, but legitimate, sequences from errors is the goal of all denoising procedures. Using paired reads also improves the applicability of the DADA2 algorithm at any step in the bioinformatic processing (not at the very beginning), thus making it a more flexible tool. As this is a requirement to perform our comparative analyses, we will use DADA2 on merged sequences, while keeping in mind that we lose stringency (retain more ESVs) by doing so.

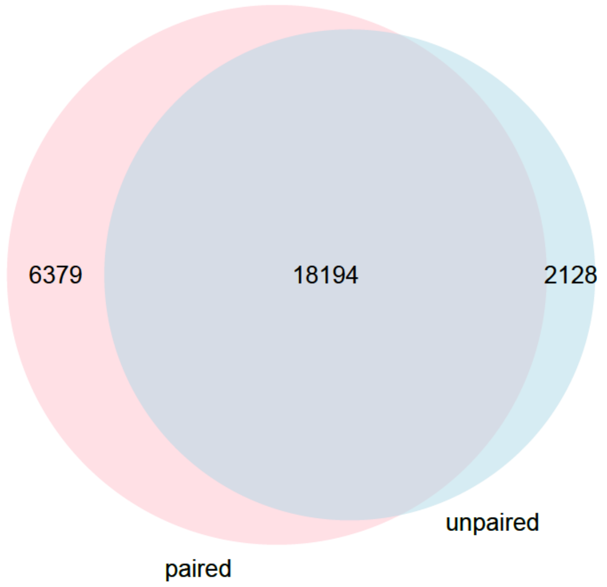


*Figure 1. Venn Diagram of the number of ESVs after applying DADA2 before (unpaired) or after (paired) merging the two reads for each sequence in the dataset analysed.*
